# Independent evolution of ab- and adaxial stomatal density enables adaptation

**DOI:** 10.1101/034355

**Authors:** Christopher D. Muir, Miquel Ángel Conesa, Jeroni Galmés

**Affiliations:** Department of Biology, Indiana University, Bloomington, IN 47405, USA; Biodiversity Research Centre and Botany Department, University of British Columbia, Vancouver, British Columbia V6T 1Z4, Canada; Research Group on Plant Biology under Mediterranean Conditions, Departament de Biologia, Universitat de les Illes Balears, Ctra. Valldemossa km 7.5, E-07122, Palma, Spain

**Keywords:** Adaptation, correlated evolution, developmental constraint, phylogenetic comparative methods, quantitative genetics, *Solanum*, stomata, stomatal ratio

## Abstract

- Are organisms free to reach their adaptive optima or constrained by hard-wired developmental programs? Recent evidence suggests that the arrangement of stomata on abaxial (lower) and adaxial (upper) leaf surfaces may be an important adaptation in plants, but stomatal traits on each surface likely share developmental pathways that could hamper evolution.
- We reviewed the quantitative genetics of stomatal density to look for loci that (1) affected ab- or adaxial density independently or (2) pleiotropically affected stomatal density on both surfaces. We also used phylogenetic comparative methods to test for independent versus correlated evolution of stomatal traits (density, size, and pore index) on each surface from 14 amphistomatous wild tomato taxa *(Solanum*; Solanaceae).
- Naturally occurring and laboratory-induced genetic variation alters stomatal density on one surface without affecting the other, indicating that development does not strongly constrain the spectrum of available mutations. Among wild tomato taxa, traits most closely related to function (stomatal pore index and density) evolved independently on each surface, whereas stomatal size was constrained by correlated evolution.
- Genetics and phylogenetics demonstrate mostly independent evolution of stomatal function on each leaf surface, facilitating largely unfettered access to fitness optima.

## Introduction

Are traits able to evolve independently of one another or they constrained by development, genetic, or functional connections? Here, we examine whether stomata on the abaxial (‘lower’) surface of the leaf evolve independently of adaxial (‘upper’) stomata. Stomata are microscopic pores on the leaf surface formed by a pair of guard cells. The density, size, and arrangement of stomata on a leaf set the maximum stomatal conductance to CO_2_ diffusing into a leaf and the amount of water that transpires from it (Parkhurst, 1978; Sack et al., 2003; Franks and Farquhar, 2001; Galmés et al., 1975). Hence, stomatal traits like density, size, and ratio of upper to lower stomata have strong effects on carbon assimilation and water-use efficiency.

An unresolved question is whether stomatal size and density on each leaf surface can evolve independently or are tethered together by shared development. Stomata are most often found only on the lower leaf surface (hypostomy), but occur on both surfaces (amphistomy) in some species (Metcalfe and Chalk, 1950; Parkhurst, 1978; Mott et al., 1984), especially herbs (Salisbury, 1927; Muir, 2015) and plants from open habitats (Mott et al., 1984; Gibson, 1996; Jordan et al., 2014). The proportion of stomata found on the upper surface also tends to increase during domestication, even as the total stomatal density stays constant (Milla et al., 2013). Amphistomy increases CO_2_ diffusion within the leaf by opening up a second parallel pathway in the intercellular airspace for diffusion from substomatal cavities to mesophyll cell walls. However, stomata on the upper surface in particular may be costly. For example, upper stomata increase the susceptibility to rust pathogens in *Populus* (McKown et al., 2014). Amphistomy may also cause the palisade mesophyll to dry out under strong vapor pressure deficits (Buckley et al., 2015). Muir (2015) reviewed the literature on other possible fitness costs.

It is tempting to explain the striking diversity in stomatal ratio as the result of natural selection optimally balancing the fitness costs and benefits. For this to be true, stomatal traits on both surfaces must be free to evolve independently. There are two reasons why independent evolution may be difficult. First, upper and lower stomata share developmental pathways, so mutations that alter the size or patterning on one surface could pleiotropically affect stomata on the other surface. Second, epidermal patterning may be tightly linked to, and therefore constrained by, overall ab-adaxial patterning in the leaf. In bifacial leaves with well differentiated spongy and palisade mesophyll layers ab-adaxial polarity is established very early in leaf development and required for blade outgrowth (Waites and Hudson, 1995; McConnell and Barton, 1998). If stomatal development is integrated into overall adaxial/abaxial patterning through shared regulatory pathways, then mutations that alter stomatal ratio could pleiotropically disrupt normal leaf development. Since spongy and palisade mesophyll layers specialize in CO_2_ diffusion and light harvesting, respectively, to optimize carbon gain, such disruption could be deleterious. Hence, populations may be unable to respond to selection on stomatal ratio because of antagonistic pleiotropy, preventing them from reaching their adaptive optima.

Multiple reviews of stomatal development conclude that stomatal traits are independently controlled on each surface (Lake et al., 2002; Bergmann and Sack, 2007), but there is little evidence for this claim. Nor is there strong evidence from the developmental literature for tight linkage between ab-adaxial polarity and stomatal development (Kidner and Timmermans, 2010; Pillitteri and Torii, 2012). To fill this gap, we use two complementary methods to directly test whether upper and lower stomatal traits can evolve independently. First, we reviewed the genetic literature for loci that effect stomatal density. If upper and lower stomatal densities can evolve independently, then we expected to find loci that specifically alter density on the upper or lower surface, but not both. Second, we took a phylogenetic comparative approach to ask whether upper and lower stomata evolve independently among a closely related group of wild tomato species *(Solanum* sect. *Lycopersicon* (Miller) Wettstein in Engler & Prantl, sect. *Lycopersicoides* (A. Child) Peralta, and sect. *Juglandifolia* (Rydberg) A. Child; Solanaceae) grown in a common garden. Both genetic and phylogenetic comparisons indicate that stomatal density on one leaf surface can evolve independently of density on the other surface. This implies that natural or artificial selection should be able to optimize the ratio of stomata on the upper and lower surface.

## Methods and Results

### Genetics reveals partially independent control of ab- and adax- ial stomatal density

We reviewed the literature on quantitative trait locus (QTL) mapping and genome-wide association studies (GWAS) of stomatal traits within and between species. We searched broadly using Google Scholar and ISI Web of Knowledge, as well as by looking through citations of and literature cited within studies we found. Seven studies of four genera *(Brassica, Populus, Solanum, Oryza*) measured separate ab- and adaxial stomatal trait loci (Table 1). Six used QTL mapping; one used GWAS. We restricted our analysis to stomatal density because not all studies measured stomatal size. We counted the number of loci that altered ab- or adaxial density, but not both (‘independent loci’) and loci that altered ab- and adaxial density in the same direction (‘shared loci’). For example, if two loci increased abaxial density and two loci increased adaxial density, and one locus for each surface colocalized, then we counted this as two independent loci (one abaxial, one adaxial) and one shared locus. If reported, we also indicated whether the authors found a significant genetic correlation between ab- and adaxial stomatal density across all genotypes. One study measured stomatal QTL at both ambient and elevated [CO_2_] (Rae et al., 2006; Ferris et al., 2002); we used only data from the ambient [CO_2_] treatment. In another study, QTL were determined at two life stages (Laza et al., 2010); we counted QTL if they affected density at one life stage or both. Finally, some studies measured QTL in the same species (Ishimaru et al., 2001; Laza et al., 2010) or even the same lines (Chitwood et al., 2013; Muir et al., 2014b), albeit under different conditions, and are clearly not independent data points.

Genetic studies reveal some correlation between stomatal densities on each surface, but in all cases there are loci which alter stomatal density on one surface independently of the other (Table 1). In some cases, there was no detectable genetic correlation between stomatal densities on each surface, which would optimally facilitate adaptive evolution. However, with few studies it is difficult to generalize about how strongly genetic covariation between stomatal traits on each surface would constrain responses to selection on microevolutionary timescales. It is also difficult to predict macroevolutionary constraints from genetic correlations within species, as genetic correlations themselves may evolve. Therefore, we next looked at macroevolutionary patterns of correlated evolution using a phylogenetic approach.

### Stomatal pore area and density, but not size, evolve independently on each surface

#### Stomatal trait measurements

We measured stomatal density (SD) and guard cell length (GCL) from ab- and adax-ial surfaces of 14 wild tomato species. Supporting Information Table S2 lists species names and Tomato Genetic Resource Center accession numbers of seed sources. There were 3–5 biological replicates per species, except the glabrous *S. chilense*, for which we could only get an accurate count from one replicate. Species were grown in a common garden at the experimental field at the University of the Balearic Islands, as described in Muir et al. (2015). We made polyvinylsiloxane (Kerr Extrude Medium, Orange, California, USA) casts of leaf surfaces from fully expanded adult leaves. We painted casts with a thin coat of nail polish and mounted this on a glass slide to count the number of stomata from three (proximal, medial, and distal) 0.571 mm^2^ portions of the leaf area unobstructed by major veins. We measured average GCL on 20 stomata per portion, 60 stomata per leaf surface examined. For each leaf surface, we calculated Stomatal Pore Index (SPI) as SD × GCL^2^, where SD and GCL are in units of stomata per mm^2^ and mm, respectively. SPI indicates what proportion of the leaf surface is occupied by stomatal pore and is closely related to maximum stomatal conductance (Sack et al., 2003). Total SD and SPI were calculated as the sum of ab- and adaxial values:

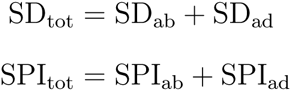

The _ab_ and _ad_ subscripts denote stomatal traits values on the ab- and adaxial surface, respectively. We measured total leaf stomatal conductance to CO_2_ (*g*_s_) under ambient CO_2_ concentrations (400 ppm) using an open-path infrared gas exchange analyzer (LI-6400 or LI-6400XT, LI-COR Inc., Lincoln, NE, USA) as described in Muir et al. (2015). Stomatal conductance was measured under optimal conditions to approach maximum *g*_s_. Steady-state measurements were taken at midday with saturating irradiance (photosynthetically active radiation set to 1500 *μ*mol quanta m^-2^ s^-1^), moderate relative humidity (40-âĂŞ60%), and 25°C leaf temperature.

#### Phylogenetic methods

Stomatal traits on each surface clearly differ from one another (Figure 1). The abaxial (lower) surface of tomato leaves usually have higher stomatal density and stomatal pore index, but smaller guard cells. Although stomata from each surface clearly differ overall, shared developmental pathways may nevertheless constrain how stomatal traits on each surface evolve. We tested whether ab- and adaxial stomatal traits evolve independently using phylogenetic comparative methods. If ab- and adaxial traits can evolve independently, then phylogenetic models assuming zero covariance between traits should outperform models with covariance. Conversely, if ab- and adaxial stomatal traits share common developmental pathways that constrain their evolution, then models with positive covariance should outperform models without covariance. To test this, we compared six models using the R package **mvMORPH** (Clavel et al., 2015). We used a maximum likelihood phylogenetic tree inferred from 18 genes (Haak et al., 2014) using RAxML version 8.1.24 (Stamatakis, 2014). All models allow separate average values for ab- and adaxial traits, but differ in two respects. First, we compared Brownian motion (BM) to Ornstein-Uhlenbeck (OU) models. In both BM and OU models, trait values evolve at rate *σ*. The only difference between BM and OU models is that the OU model includes an extra parameter (denoted *α*) that pulls species back faster toward the long-run average (denoted *θ*). Note that BM versus OU comparison only tests how tightly stomatal traits are constrained to evolve around the long-run average, not whether ab- and adaxial stomata evolve independently. We tested for independent evolution by comparing BM and OU models with and without evolutionary covariance between leaf surfaces. We compared two BM models for each trait, one in which Cov(*σ*_ab_, *σ*_ad_) is estimated (‘covary’ model) and another with the constraint Cov(*σ*_ab_, *σ*_ad_) = 0 (‘independent’ model). Similarly, we competed four OU models for each trait, three that allowed covariance between ab- and adaxial evolution and one in which they evolved independently. Specifically, we tested for covariance between diffusion rates (Cov(*σ*_ab_, *σ*_ad_) estimated, Cov(*α*_ab_, *α*_ad_) = 0), covariance between return rates (Cov(*α*_ab_, *α*_ad_) estimated, Cov(*σ*_ab_, *σ*_ad_) = 0), or covariance between both diffusion and return rates (Cov(*σ*_ab_, *σ*_ad_) and Cov(*α*_ab_, *α*_ad_) estimated). The ‘independent’ model constrained both Cov(*σ*_ab_, *σ*_ad_) = 0 and Cov(*α*_ab_, *α*_ad_) = 0. We incorporated measurement error using the standard error across biological replicates within species (Pennell et al., 2015). Because we could not estimate measurement error for *S. chilense*, we used the average measurement from the other species instead. All traits were log-transformed for normality. We compared model fit using Akaike Information Criterion corrected for small sample size (AICc).

**Fig. 1.**
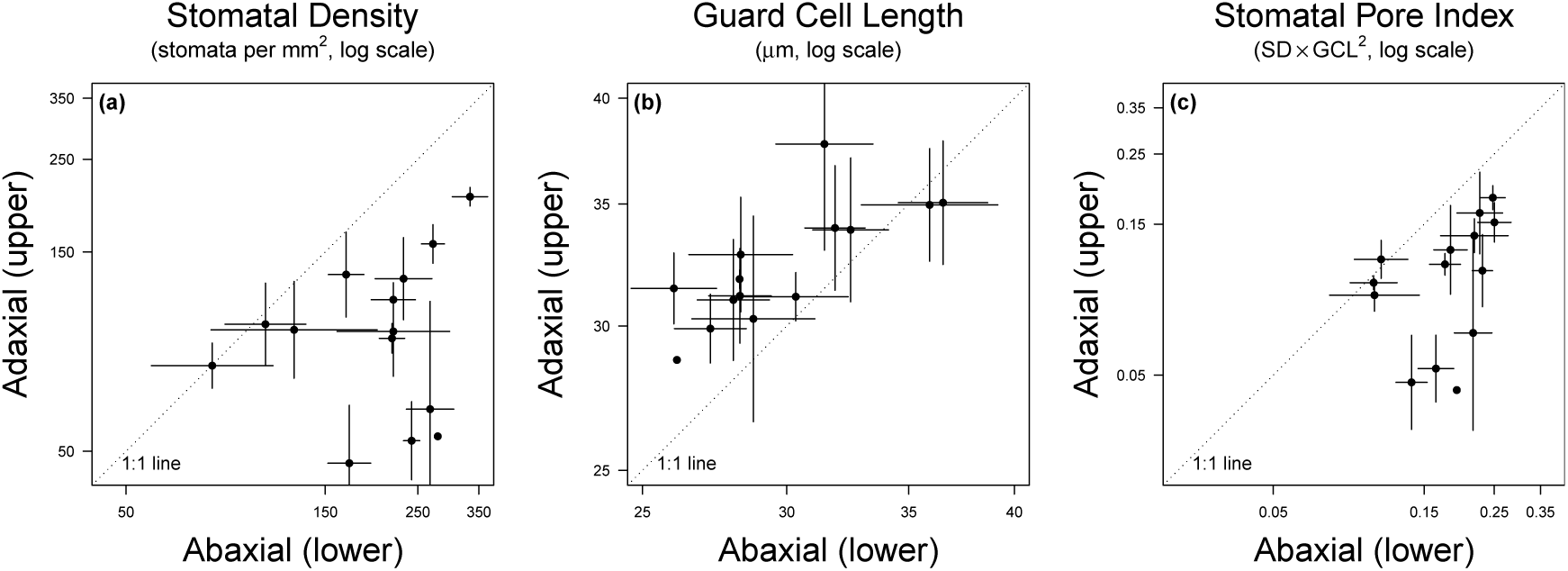
Ab- and adaxial stomatal density (SD; panel **(a)**) and stomatal pore index (SPI; panel **(c)**) evolve independently, whereas the guard cell lengths (GCL; panel **(b)**), a measure of stomatal size, positively covaries over evolution. In horizontally-oriented tomato leaves, ab- and adaxial surfaces are the lower and upper surface, respectively. Adaxial SD and SPI values tend to be lower than abaxial ones (most points fall below 1:1 line), whereas adaxial stomata tend to be larger (higher GCL) than abaxial ones. Each point is mean trait value for one of 14 wild tomato species; lines are +/- one standard deviation. One species, *S. chilense*, was only sampled once and therefore the standard deviation could not be estimated.

SD and SPI evolution are constrained but ab- and adaxial traits are uncorrelated. For both traits, OU models fit better than BM models, and models with covariance between leaf surfaces performed worse than those without covariance (Table S1). We found the opposite pattern for GCL. Evolution of this trait was best described by a model without constraint but including covariance between ab- and adaxial GCL. Since SPI_tot_ is closely related to stomatal conductance in these species (Figure 2), independent evolution of SPI_tot_ suggests little evolutionary constraint on how stomatal conductance is partitioned across surfaces in different species.

**Fig. 2.**
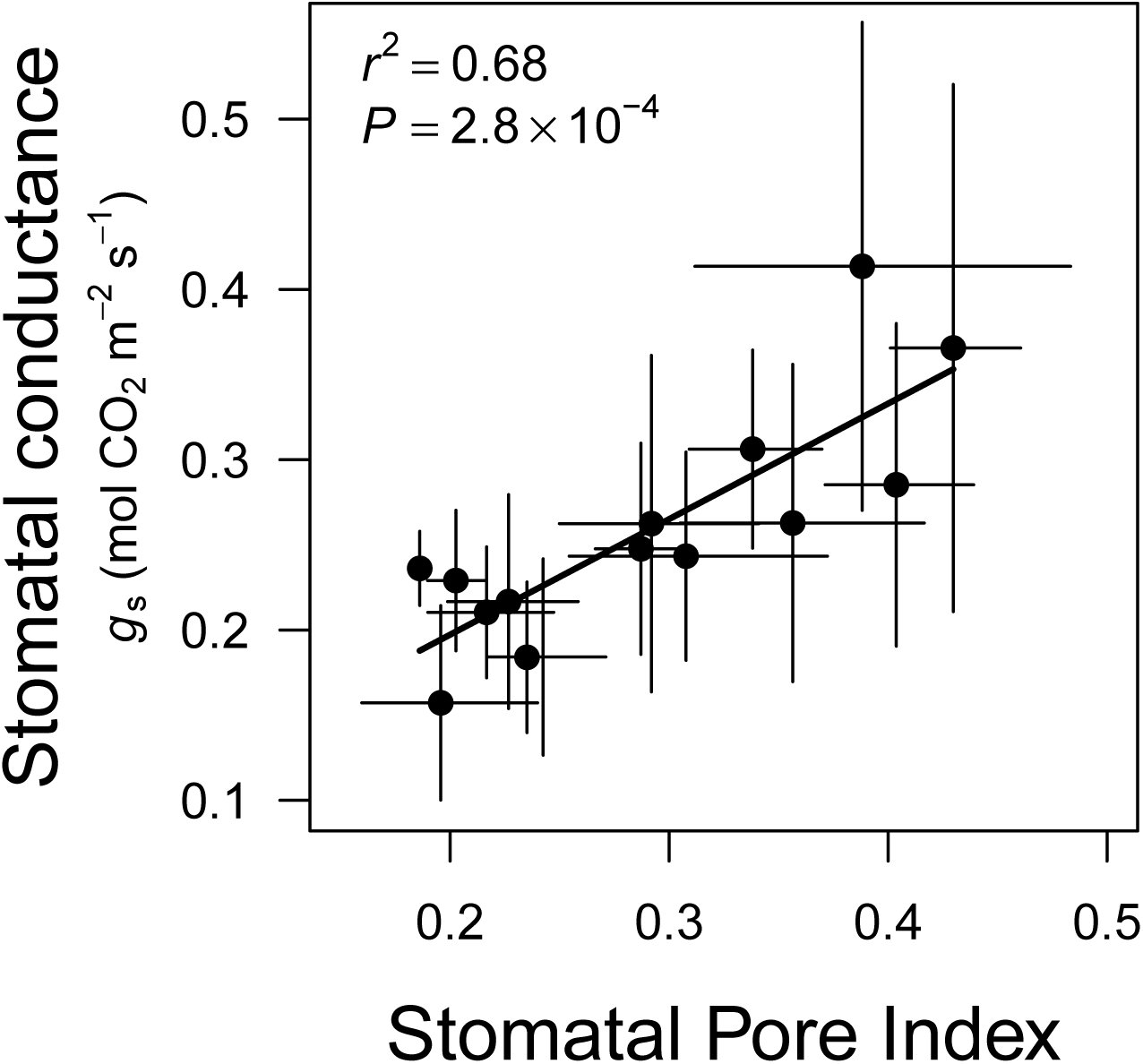
Stomatal conductance to CO_2_ (*g*_s_) is directly proportional to stomatal pore index (SPI) in wild tomato species. *g*_s_ was measured at ambient CO_2_ concentrations (400 *μ*mol CO_2_ mol^-1^ air), saturating irradiance (1500 *μ*mol quanta m^-2^ s^-1^), 25°C leaf temperature, and 40–60% relative humdity. Each point is the species mean; error bars are +/- one standard deviation.

## Discussion

Adaptive evolution may be constrained if traits cannot evolve independently. In particular, if traits share developmental pathways, then they may be unable to respond differentially to selection. In this study, we examined whether stomata on the abaxial (lower) and adaxial (upper) surfaces can evolve independently. We adduce two new lines of evidence which suggest that stomatal function on each surface can readily respond to selection. First, species possess heritable variation that allows partially independent evolution of stomatal densities in response to selection; every study reviewed found loci which alter stomatal density on one surface but not the other. Second, the anatomical trait most closely connected to stomatal conductance, stomatal pore index (Sack et al., 2003), evolves independently on ab- and adaxial surfaces among wild tomato species. Together, these new lines of evidence demonstrate that natural selection on stomatal arrangement is not strongly constrained by development, although we lacked statistical power to detect weak constraint. It is therefore likely that variation in how stomatal conductance is partitioned between leaf surfaces is due to adaptive rather than nonadaptive forces.

Indeed, much recent evidence indicates that selection finely tunes the ratio of stomata on the upper and lower leaf surface, although the adaptive significance of variation in stomatal ratio is unresolved. Stomatal ratio affects leaf function, increasing CO2 diffusion (Parkhurst, 1978; Parkhurst and Mott, 1990; Gutschick, 1984; Parkhurst, 1994) and hydraulic conductance outside the xylem (Buckley et al., 2015). As predicted, amphistomy seems to be more common in circumstances when efficient CO2 supply is important, such as high irradiance (Mott et al., 1984; Gibson, 1996; Smith et al., 1997; Jordan et al., 2014), thick leaves (Parkhurst, 1978; Muir et al., 2014a), herbaceous growth form (Salisbury, 1927; Muir, 2015), and domestication (Milla et al., 2013). Despite potential benefits of amphistomy, most plant species are hypostomatous, implying a fitness cost of upper stomata, such as increased infection by foliar pathogens (Gutschick, 1984; McKown et al., 2014). For example, ‘upside-down’ (resupinate) leaves with the abaxial surface facing upward have re-evolved hypostomy (Lyshede, 2002), strongly implying a cost of upward facing stomata.

To optimally balance fitness costs and benefits, natural selection must be able to change stomatal traits on one surface independently of the other. The present study shows that this is likely true and strikingly consistent on micro- and macroevolutionary timescales. Among *Populus trichocarpa* populations and *Solanum* species, the ratio of adaxial to abaxial SPI (SPI ratio) evolves mostly by changes in stomatal density rather than guard cell size. Within *Populus*, populations are more amphistomatous at Northern latitudes with shorter growing seasons that may select for faster carbon assimilation (McKown et al., 2014; Kaluthota et al., 2015). Latitudinal variation *Populus trichocarpa* is due mostly to adaptive variation in adaxial stomatal density (McKown et al., 2014; Porth et al., 2015). Stomatal density rather than size may have responded more readily to selection because there is no genetic covariance between ab- and adaxial stomatal density, permitting independent evolution (Porth et al., 2015). In contrast ab- and adaxial guard cell length positively covary, likely constraining evolution. Similarly, we found that over macroevolutionary timescales most of the variation in SPI among wild tomato species is due to changes in adaxial stomatal density rather than size. Indeed, stomatal density on each surface evolved independently, whereas guard cell lengths positively covaried (Table S1). Adaptive evolution will likely take advantage of traits that evolve independently because this minimizes antagonistic pleiotropy. In a previous study, we found that loci affecting adaxial stomatal density were likely fixed by selection, but we did not measure stomatal size (Muir et al., 2014b). Overall, patterns within and between species indicate that selection on SPI ratio leads to greater change in stomatal densities rather sizes on each surface. Based on the analysis here, we conclude that changing stomatal density on one surface incurs less cost than changing size because the former is less constrained by shared developmental pathways.

We caution that there are limitations of our analysis. First, although some loci alter stomatal traits on one surface independently of the other, there are also loci that affect both surfaces, leading to significant genetic correlations in some species (Table 1). Such genetic correlations will slow adaptation even if they do not prevent populations from eventually reaching an adaptive optimum in the long run. For example, the relatively high genetic correlation between ab- and adaxial stomatal density in *Oryza* may contribute to low variation in stomatal ratio between species of this genus (Giuliani et al., 2013). Second, the sample size of the phylogenetic comparisons is small and thus not statistically powerful. However, simulations show that model identification (e.g. Brownian motion versus Ornstein-Uhlenbeck) is usually correct, even when sample sizes are moderate (Cressler et al., 2015; Ho and Ané, 2014). The dataset was also powerful enough to find significant correlated evolution of guard cell size on ab- and adaxial surfaces, which we interpret as evidence of shared developmental pathways. However, we cannot rule out some level of correlated evolution for stomatal density and pore index below our threshold to detect. Finally, stomatal traits measured in a common garden may be different than what occurs naturally. For example, the ratio of stomatal density and size changes in response to light (Gay and Hurd, 1975) and water stress (Galmés et al., 1975). Despite *ad libitum* watering and fertilizer, our common garden in a Mediterranean climate may have been more stressful for some tomato species than others, depending on their habitat of origin, perhaps inducing stress-response phenotypes.

**Table 1.** Many loci alter ab- or adaxial stomatal density independently, while others affect both surfaces. We reviewed seven studies (key to reference numbers below) in four genera. Loci were identified using quantitative trail locus mapping (QTL) or genome-wide association studies (GWAS). Shared loci altered both ab- and adaxial stomata density, whereas independent loci affected one or the other. The Data Source column refers to the table or figure in the reference where we found data. Muir et al. (2014b) did not report these analyses, but we calculated number of QTL using the same methods. We also indicate whether the study reported significant genetic correlation between ab- and adaxial stomatal density across all genotypes.

| Reference | Taxa | Method | Shared loci | Independent loci | Data source | Significant genetic correlation? |
| --- | --- | --- | --- | --- | --- | --- |
| (1) | *Brassica oleracea* L. | QTL | 2 | 1 | Table 1 | not reported |
| (2, 3) | *Populus trichocarpa* Torr. & A. Gray ex Hook. | GWAS | 0 | 9 | Table 3<sup>2</sup> | no |
| (4) | *Solanum* species | QTL | 7 | 17 | unpub. result | no |
| (5) | *Solanum* species | QTL | 7 | 14 | Supplemental Figure 10 | yes |
| (6, 7) | *Populus* species | QTL | 0 | 6 | Figure 4<sup>7</sup> | not reported |
| (8) | *Oryza sativa* L. | QTL | 1 | 2 | Table 1 | yes |
| (9) | *Oryza sativa* L. | QTL | 1 | 8 | Table 3 | yes |

<sup>1</sup> Hall et al. (2005); <sup>2</sup> McKown et al. (2014); <sup>3</sup> Porth et al. (2015); <sup>4</sup> Muir et al. (2014b); <sup>5</sup> Chitwood et al. (2013); <sup>6</sup> Ferris et al. (2002); <sup>7</sup> Rae et al. (2006); <sup>8</sup> Ishimaru et al. (2001); <sup>9</sup> Laza et al. (2010)

**Table 2.** Phylogenetic comparisons reveal independent evolution of ab- and adaxial stomatal density (SD) and stomatal pore index (SPI), but shared developmental pathways for ab- and adaxial guard cell length (GCL). We compared Brownian motion (BM) and Ornstein-Uhlenbeck (OU) models. Under the BM model, average trait values (*θ*) evolve without bounds at rate *σ*, whereas under the OU model, trait values are bounded. *α* is the return rate toward *θ* in the OU model. For both OU and BM models, we compared models with (‘covary’) and without (‘independent’) covariance between ab- and adaxial traits. We compared models using Akaike Information Criteria corrected for small sample size (AICc). ΔAICc for a model is the difference its AICc and that of the model with lowest AICc. Hence, for the best-supported model ΔAICc = 0. *k* is the number of parameters estimated for a particular model.

| Model | Model parameters | | | $k$ | $\Delta\text{AICc}$ | | |
| --- | --- | --- | --- | --- | --- | --- | --- |
| | $\theta$ | $\sigma$ | $\alpha$ | | GCL | SD | SPI |
| BM1 | separate | covary | – | 5 | 0 | 6.5 | 7 |
| BM2 |  | independent | – | 4 | 10.1 | 11.8 | 12.4 |
| OU1 |  | covary | covary | 8 | 19.9 | 13.4 | 14.9 |
| OU2 |  | covary | independent | 7 | 7.8 | 7.5 | 5.5 |
| OU3 |  | independent | covary | 7 | 7.8 | 3.3 | 4.6 |
| OU4 |  | independent | independent | 6 | 13.1 | 0 | 0 |

We recommend that future genetic and comparative studies of stomatal traits report separate ab- and adaxial values for stomatal density and size. We also need to determine developmental connections between abaxial/adaxial pattern specification and epidermal development. For example, SPCH SILENT, an *Arabidopsis* mutant that relatively normal adaxial stomatal density but no abaxial stomata (Dow et al., 2014), suggesting possible links between SPCH and abaxial/adaxial patterning. The molecular mechanisms may explain how stomata often develop differently on each surface and why asymmetry between surfaces readily evolves.

## Acknowledgements

CDM was supported by an Evo-Devo-Eco Network (EDEN) research exchange (NSF IOS #0955517). The research was supported by project AGL2013–42364-R (Plan Nacional, Spain) and UIB Grant 15/2105 awarded to JG. We acknowledge Miquel Truyols and collaborators of the UIB Experimental Field and Greenhouses for their technical support. Matt Pennell provided feedback.

## Author contribution statement

CDM, MAC, and JG designed and carried out the experiment. CDM analyzed the data and wrote the manuscript.

## Supporting Information

**Table S1.** Wild tomato species and Tomato Genetic Resource Center (TGRC) accession numbers.

| Species | TGRC Number |
| --- | --- |
| *S. arcanum* Peralta | LA2153 |
| *S. cheesmaniae* (Riley) Fosberg | LA1035 |
| *S. chilense* Dunal | LA1782 |
| *S. chmielewskii* D.M.Spooner, G.J.Anderson & R.K.Jansen | LA1327 |
| *S. galapagense* S.C. Darwin & Peralta | LA0930 |
| *S. habrochaites* S. Knapp & D.M. Spooner | LA2196 |
| *S. lycopersicoides* Dunal | LA2951 |
| *S. lycopersicum* var. *cerasiforme* (Dunal) D.M. Spooner, G.J. Anderson & R.K. Jansen | LA1320 |
| *S. neorickii* D.M. Spooner, G.J. Anderson & R.K. Jansen | LA1322 |
| *S. pennellii* Correll | LA1272 |
| *S. pennellii* var. *puberulum* Correll | LA1926 |
| *S. peruvianum* L. | LA2964 |
| *S. pimpinellifolium* L. | LA0114 |
| *S. sitiens* I.M. Johnst. | LA4115 |

